# Long weekend sleep is linked to stronger academic performance in male but not female pharmacy students

**DOI:** 10.1101/2020.01.15.907907

**Authors:** Rehana Khan Leak, Susan L. Weiner, Manisha N. Chandwani, Diane C. Rhodes

## Abstract

**Introduction:** Poor sleep hygiene portends loss of physical and mental stamina. Therefore, maintaining a regular sleep/wake schedule on both weekdays and weekends is highly recommended. However, this advice runs contrary to the habits of university students, who may only experience recovery sleep if they sleep late on weekends. Pharmacy students at Duquesne University sit for frequent examinations, typically commencing at 7:30 AM, and they complain about fatigue. Here, we tested the central hypothesis that longer sleep durations on weekdays and weekends are linked to stronger academic performance in Pharmacy students.

**Methods:** Students in their first professional year were administered three surveys to collect data on sleep habits and factors that might influence sleep, such as roommates, long commute times, and sleep interruptions. GPAs were collected from the Dean’s office, with permission from the students.

**Results:** Longer weekend—but not weekday—sleep durations were significantly correlated with higher cumulative GPAs in men and not in women. Women achieved slightly higher cumulative GPAs than men. Students who fell asleep within 15 minutes of going to bed had higher professional-phase GPAs than those who fell asleep after an hour or more.

**Conclusion:** Our observations cannot establish causal links, but, given the body of prior evidence on the salutary properties of sleep, men in this cohort may reap benefit from recovery sleep on weekends. Rather than recommending that students force themselves awake early on weekends in an attempt to maintain a consistent sleep routine, the real-life habits of students should be given consideration.

## Introduction

> *“Sleep … Balm of hurt minds … Chief nourisher in life’s feast.”*
>
> Macbeth (2.2.46-51), by William Shakespeare

Although the sleep phase of the rest/activity rhythm increases the risk of falling vulnerable to predation, natural selection has strongly favored a consolidated period of sleep in mammals. The human rest/activity rhythm is normally entrained to the environmental photoperiod, but it can also drift in response to lifestyle factors, leading to loss of sleep or increased rebound sleep. Aside from adverse effects on metabolism, blood pressure, endocrine function, and numerous other physiological processes,^1-3^ sleep loss and irregular sleep patterns are linked to inferior cognitive measures, such as poor memory consolidation and subpar academic performance.^4-6^

There are several physiological explanations for the potential impact of sleep on academic performance. First, the glymphatic system of the brain performs its janitorial duties and clears accumulated metabolites via the cerebrospinal fluid better during the sleep phase than the activity phase.^7-9^ Second, sleep loss-induced attentional deficits are preceded by electrophysiological lapses in neuronal function, and the association between sleep loss and cognitive impairment is thought to be causal.^10-16^ Sleep has been reported to promote the consolidation of newly acquired memories, perhaps by modifying the strength of synaptic connections, including weakening synapses that were previously inactive.^17, 18^

The physiological consequences of sleep deprivation have implications on our society, including for health professionals and the patients that depend on their care. Physicians deprived of sleep for 30 hours may suffer a reduction in clinical performance by >1.5 standard deviations.^19^ Even partial sleep deprivation across sufficient days can reduce cognitive function to levels observed after total acute sleep loss.^20^ Healthcare professionals must also acquire and retain information during their professional training phase. Based on their 2008 survey on sleep hygiene awareness in student pharmacists, however, Ang and colleagues reported “a need to improve practicing pharmacists’ as well as undergraduate students’ knowledge of sleep health.”^21^ A recent study by Zeek and colleagues employed anonymous self-reporting to collect evidence that ∼50% of student pharmacists experienced suboptimal sleep durations (<7 hours), and reported being sleepy during the day.^22^ As expected, shorter sleep durations were linked to lower self-reported grade point averages, with a grade reduction for every hour of sleep lost.^22^

Cates and colleagues gathered self-reported, anonymous grade-point averages from student pharmacists and administered them the Pittsburgh Sleep Quality Index.^23^ The latter authors also discovered widespread poor sleep quality (at a level similar to postpartum mothers with infants), especially for students with lower self-reported GPAs. In the same study, female pharmacy students reported less difficulty with maintaining sleep durations than male students.^23^ Recent findings suggest that women display superior sleep measures, such as higher sleep quality and longer sleep durations, but women have also reported poorer quality sleep, including more insomnia, compared to men.^24, 25^ Thus, the complex interactions between biological sex and sleep metrics need to be examined further.

In the Pharmacy program at Duquesne University, exams are administered at 7:30 AM in the morning (6:30 or 7:00 AM for special needs students), and classes commence at 8:00 AM. Despite the need for early-morning awakenings, anecdotal comments from Pharmacy students suggest that they often stay up late at night, cramming for the 7:30 AM examinations, and then “crash” on weekends by oversleeping. Experts have recommended a consistent sleep schedule as the best practice for good sleep hygiene and superior academic performance, as was recently rigorously demonstrated by Okano and colleagues in students enrolled at Massachusetts Institute of Technology,^4^ and by Phillips and colleagues in students enrolled at Harvard University.^26^ The data collected in both of these studies support the view that regular sleep habits improve student grades. Unfortunately, the abovementioned studies did not present data stratified by weekday versus weekend sleep. Therefore, we investigated the potential impact of long weekend versus weekday sleep durations on grade point averages (GPAs) of pharmacy students at Duquesne University. GPAs were acquired from the Dean’s office, rather than relying on self-reporting. Gender was added as an independent variable in our analyses.

## Methods

### Study Design

Ethics approval for three surveys was granted by the Institutional Review Board at Duquesne University, and their standardized consent forms were employed. All three surveys were administered to the same students. First, a mandatory online homework assignment (Survey 1) was administered in the Ability Based Laboratory Experience (ABLE) course at Duquesne University. Students register for this course in the second semester of Year 1 of the four-year professional phase, after completion of the two-year preprofessional phase. Out of 152 enrolled students, all completed the daily online Survey 1 (**Appendix 1**) to record bedtimes, sleep times, and awakenings for three consecutive weeks mid-semester.

Survey 2 was voluntary. In Survey 2, demographic information and specific permission to publish the data from Survey 1 (on a separate page from demographic data) were collected from 125 students (**Appendix 1**). If the consent form on the separate page was left unsigned, the entire packet from that student and their Survey 1 data were not used. Even if the consent from was signed by the student, it was nonetheless detached from Survey 2 to maintain anonymity, not allowing us to connect *individual* data points across Surveys 1 and 2. Rather, the data in Survey 2, including the self-reported GPAs, were used to guide the questions in the final, third survey. In other words, data from Survey 2 were used independently from surveys 1 and 3, as we were not able to link it back to the individual.

The third survey was also voluntary and deployed two months later to the same class of students, to continue to assess lifestyle factors hypothesized to impact sleep quality and academic performance, such as participation on an athletic team, nap frequency and duration, hours spent working at a job, *etc*. (**Appendix 1**). One-hundred and twenty-five students participated in Survey 3. In Surveys 2 and 3, specific permission was collected to acquire grade-point averages (GPAs) from the Dean’s office, but students could refuse to have their data analyzed and published at any time. Once data were connected across Surveys 1 and 3, and the students’ cumulative and professional-phase GPAs were sent to us by the Dean’s office, all subjects were deidentified and their names replaced with ID numbers. These third-year students were in their second semester of the professional phase (*i.e.*, the third year of the entire curriculum), and, therefore, the professional-phase GPA only consisted of data from the previous semester.

### Statistics

Data were analyzed in GraphPad Prism (Prism 8 for MacOS) and subjected to Prism’s default tests for heteroscedasticity (Bartlett’s, Brown-Forsythe, and Spearman’s test) and normality (Anderson-Darling, D’Agostino-Pearson omnibus, Shapiro-Wilk, and Kolmogorov-Smirnoff tests). When parametric assumptions were met, Pearson correlations, Student *t* tests, or ANOVAs were performed on data sets. Bonferroni *post hoc* tests were used for multiple comparisons after the appropriate ANOVA. For non-Gaussian data sets, the Kruskal-Wallis test was followed by the Dunn’s *post hoc* correction for multiple comparisons. Alpha was set at 0.05 (two-tailed).

### Inclusion/Exclusion Criteria

Data were included in the analyses and graphs only if the student granted permission. Data were excluded from the survey if the student did not grant permission. If a student granted permission but failed to complete a specific part of the survey (*i.e.*, some students did not answer every single question on each survey), the remaining data were still included in the analyses. The number of students per group were added to every figure. No outliers were removed.

## Results

The majority of participants were women (82 women / 125 total responders), 22 years of age (42 were 21 years old or younger, 67 were 22 years old, and 12 were 23-25 years old) and were not part of an athletic team (11 males and 5 females were part of athletic teams). Fourteen out of 43 male responders and 43 out of 82 female responders lived on campus and did not commute to classes. Not all demographic questions were answered by every student. Other than self-reported gender, these factors did not significantly influence sleep measures or GPA, as discussed further below.

A frequency histogram of cumulative GPAs from students who granted permission is illustrated in **Figure 1A**. Weekend sleep duration was significantly associated with cumulative GPAs collected from the Dean’s office (**Figure 1B**; one-way ANOVA; F(4, 107) = 2.621; *p* = 0.0389; passed heteroscedasticity and normality tests). Students who slept 10 or more hours per weekend night had significantly higher cumulative GPAs than students who slept 6 hours per weekend night. Women had slightly higher cumulative GPAs than men (**Figure 1C**; two-tailed *t* test; *t* = 2.418; df = 118; *p* = 0.0171; passed heteroscedasticity and normality tests). Hence, the impacts of gender and weekend sleep duration on cumulative GPAs were analyzed by two-way ANOVA (**Figure 1D**; passed heteroscedasticity and normality tests). A significant interaction between gender and hours of sleep on the weekend was observed (*p* = 0.0235, F(4, 102) = 2.954), as well as a significant effect of weekend sleep duration (*p* = 0.0059; F(4, 102) = 3.851). However, Bonferroni *post hoc* comparisons revealed that the potential impacts of longer weekend sleep durations were observed in men—and not women (**Figure 1D**). Therefore, correlation analyses between weekend sleep and cumulative GPAs were plotted separately for men and women. These latter analyses confirmed a significant correlation between weekend sleep duration and cumulative GPAs for men, but not women (**Figure 2A-B**; passed normality tests). Weekday sleep duration was not associated with cumulative GPAs in men or women (**Figure 2C-D**). Furthermore, there was a notable lack of significant correlation between the duration of sleep during the week and the duration of sleep during the weekend (**Figure 2E**; passed normality tests). Given the latter finding, those who slept longer on the weekend were therefore not sleeping either more or less during the week.

**Figure 1.**
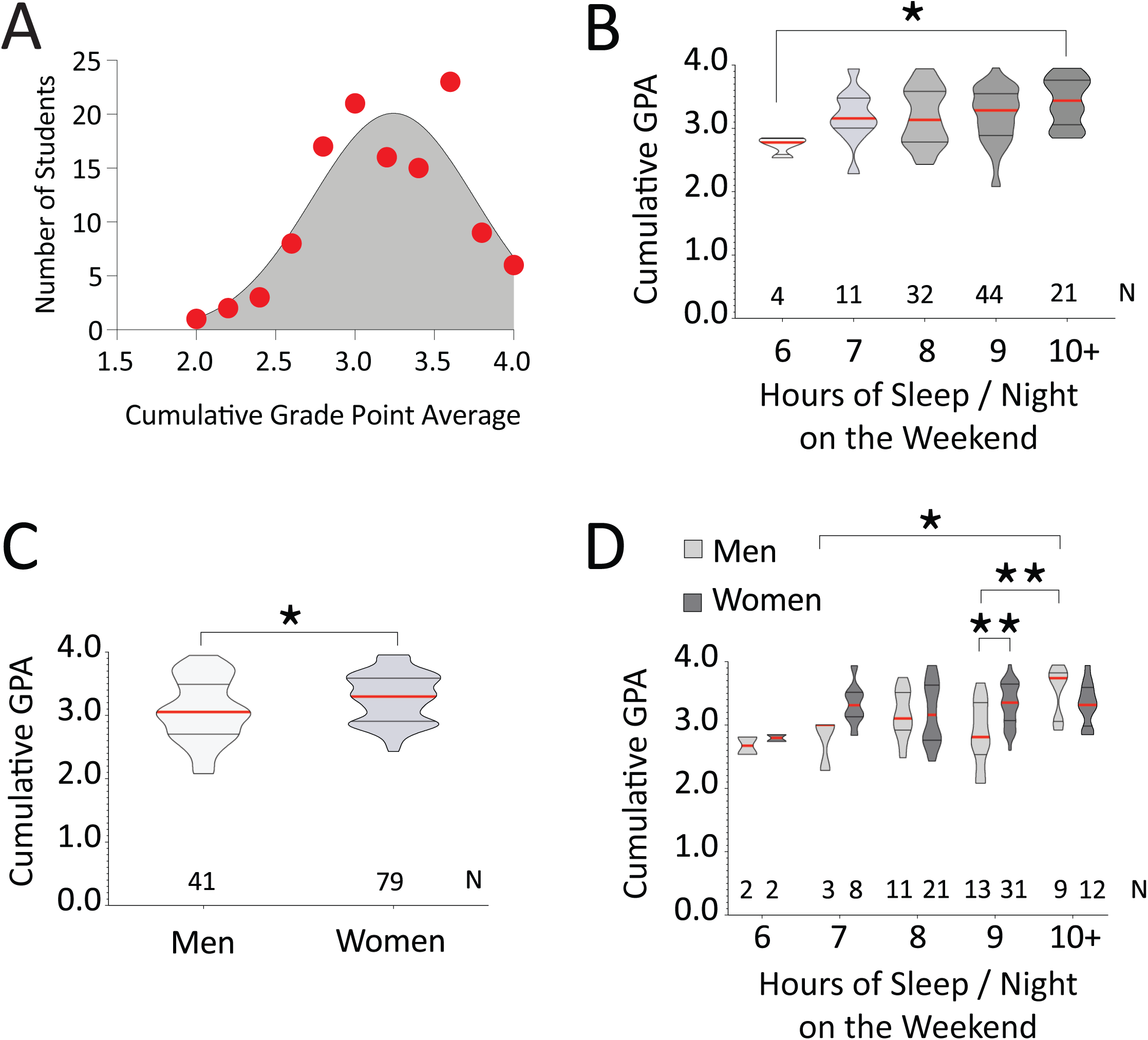
Longer durations of weekend sleep are associated with higher GPAs in pharmacy students. Students in the first year of the professional-phase pharmacy program at Duquesne University were asked to closely monitor their sleep schedules, on a daily basis, for three weeks. (**A**) Frequency histogram of cumulative GPAs collected from the Dean’s office, with signed permission from the students. Red dots indicate the raw numbers of students who achieved the cumulative GPAs shown on the Y axis. The gray fitted curve is a Gaussian least squares fitted line, according to the equation Y=Amplitude*exp(−0.5*((X-Mean)/SD)^2). **(B**) Hours of sleep on weekends were plotted against cumulative GPAs. Data are illustrated as violin plots, in which the width of the plot varies as a function of the relative density of the data points, but only within each group. Therefore, the number of participants per group is also listed above the X axis. Horizontal lines denote the quartiles of the data distribution for each group from 0-100, and each red horizontal line denotes the median. * *p* < 0.05 for groups indicated by bracketed lines, according to one-way ANOVA and Bonferroni pairwise comparisons. (**C**) Women had slightly higher cumulative GPAs than men. * *p* < 0.05 according to unpaired, two-tailed *t* test. (**D**) Violin plots of cumulative GPA as a function of gender and hours of sleep per night on weekends. * *p* < 0.05 and ** *p <* 0.01 for groups indicated by bracketed lines, according to two-way ANOVA and Bonferroni pairwise comparisons.

**Figure 2.**
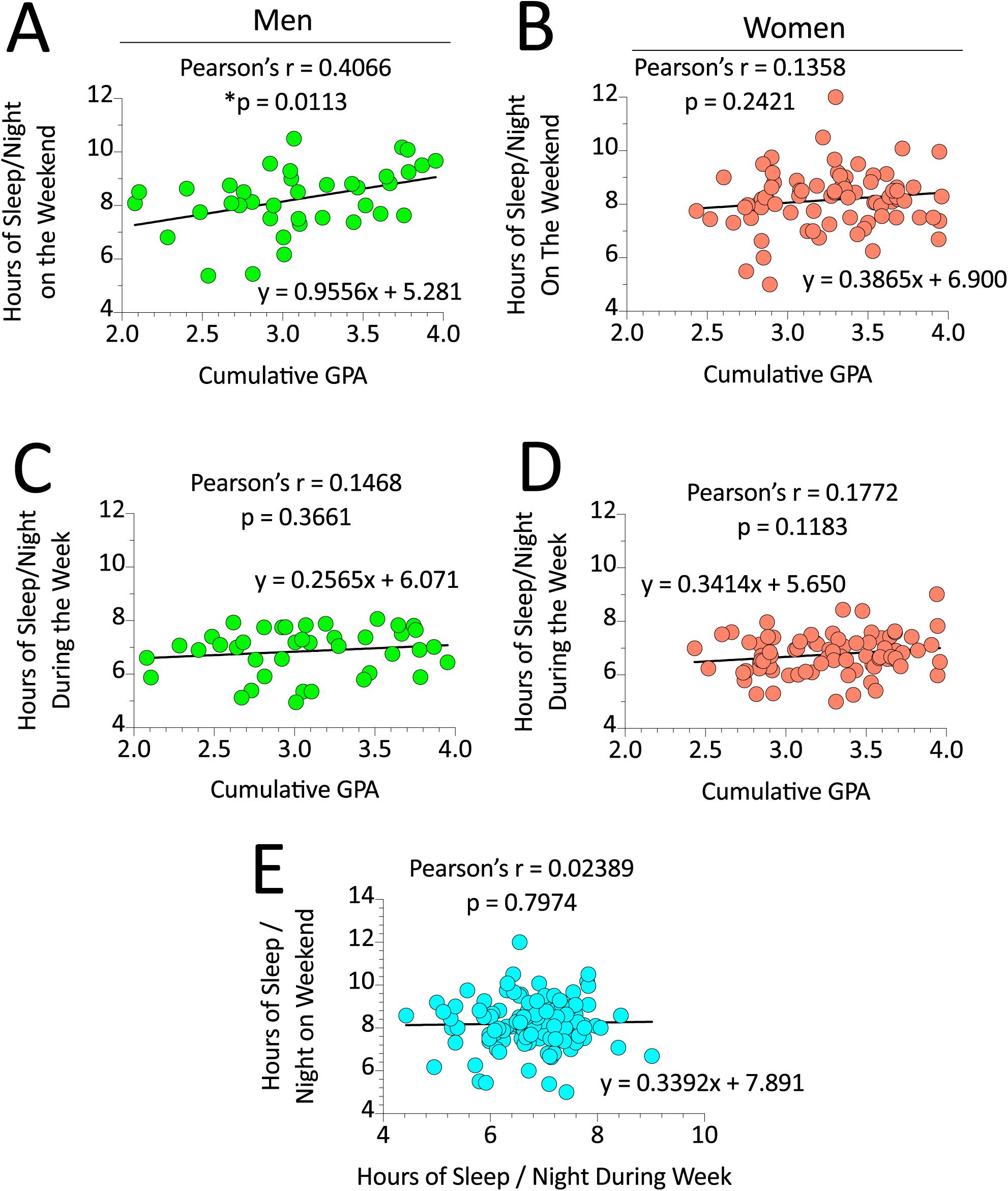
Correlation of weekend sleep durations and cumulative GPAs in male, but not female pharmacy students. (**A-B**) Pearson correlation of cumulative GPAs with hours of sleep per weekend night in men or women. (**C-D**) Pearson correlation of cumulative GPAs with hours of sleep per weekday night in men or women. (**E**) No correlation was observed between weekend and weekday sleep durations. The Pearson correlation coefficient (*r*), the *p* value to determine if the slope of the regression line is significantly nonzero, and the equation for the regression line (y = mx + b) are shown for each dataset.

Gender did not significantly influence the number of awakenings, sleep duration, or latency to fall asleep on the weekend or weekday (not shown). The average standard deviation in sleep duration for each student across the survey period (adapted from Okano *et al.* as “inconsistency in sleep duration from day to day”^4^) did not differ between men and women (**Figure 3A**; passed heteroscedasticity but failed normality tests; Mann-Whitney U statistic 1588; two-tailed *p* = 0.7788). Sleep inconsistency was not also impacted by gender when the weekend and weekday data were analyzed separately, and it was not correlated with cumulative GPA (not shown).

**Figure 3.**
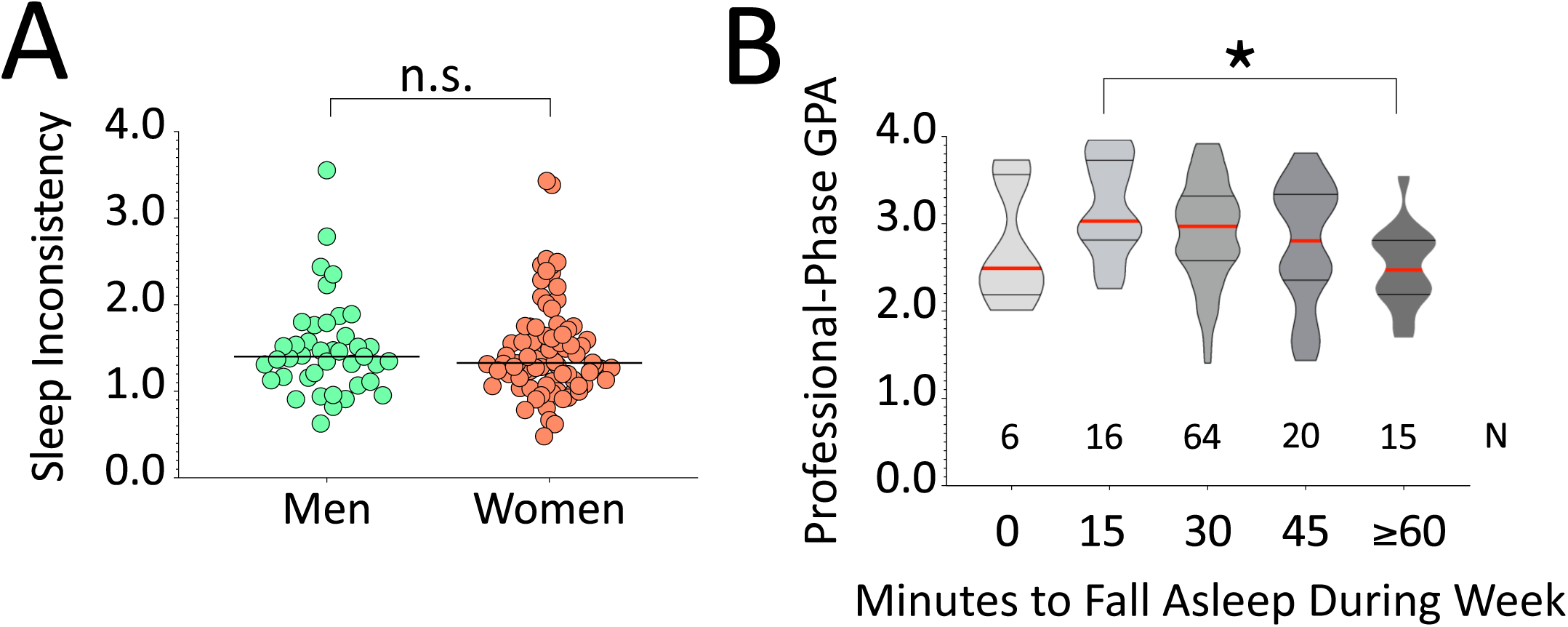
Long latencies (1h or more) to fall asleep are associated with lower GPAs. (**A**) Scatterplots of sleep inconsistency, defined as the average standard deviations of sleep duration per student across three weeks, as a function of gender. n.s. = not significantly different according to two-tailed Mann-Whitney U test. (**B**) Violin plots of professional-phase GPA (third-year GPAs only), as a function of the number of minutes to fall asleep on weekdays. * *p* < 0.05 for groups indicated by bracketed lines, according to one-way ANOVA and Bonferroni pairwise comparisons.

Other notable measures were not statistically significantly related to cumulative GPAs, including the number and duration of naps, sleep interruptions, commuting, and the number of hours of job-related work per week. Cumulative GPAs were also not significantly associated with commute duration. One exception was that the number of minutes to fall asleep after entering bed was significantly associated with GPAs from the professional phase, in an inverted U-shaped pattern (**Figure 3B**; one-way ANOVA; F(4, 116) = 2.763; *p* = 0.0308; passed heteroscedasticity and normality tests). Subjects who fell asleep within 15 minutes, on average, had significantly higher professional-phase GPAs than those who needed one or more hours. This advantage, however, was not observed in those who reported falling asleep immediately upon entering bed.

## Discussion

The main novel finding of the present study is that weekend sleep duration explained a significant proportion of the variance in the cumulative GPAs of male, but not female student pharmacists. Our students also diverge from previous studies in that we failed to observe a correlation between academic performance and weekday sleep duration,^4, 22^ perhaps due to early exam schedules, combined with a high frequency of assignments and exams (four exams/semester for multiple courses). Given the lack of significant correlations between academic performance and weekday sleep durations (including the night before an exam), our central hypothesis was only partially supported.

Given the beneficial impact of sleep on cognitive function, we had expected to find a relationship between sleep duration and cumulative GPAs in men as well as women. We did not observe higher sleep inconsistency in men compared to women, in contrast to the work of Okano and colleagues.^4^ Thus, gender differences in academic performance in our student cohort are not readily explained by the latter measure. It should be noted that differences across sleep studies in academia are not unexpected, given the unique demographics of students enrolled at different institutions. These discrepancies highlight the importance of continued work on this topic.

The second main finding of the present study is the U-shaped graph of professional-phase GPAs plotted as a function of the reported time to fall asleep. Taking one hour or more to fall asleep was associated with lower professional-phase GPAs than for those who required, on average, 15 minutes. Those who fell asleep as soon as their heads hit the pillow enjoyed no such advantage, perhaps because they were too exhausted from sleep deprivation, which, as argued in the Introduction, is associated with poor academic performance. These observations suggest that additional information on sleep-delaying factors, such as blue light exposure from electronic devices and anxiety-related insomnia, should be investigated in this cohort, particularly during the professional phase.

Given our results, we speculate that men may be more vulnerable than women to the negative sequelae of foreshortened weekend sleep and may benefit from sleeping longer on the weekend, although it seems reasonable to recommend that both sexes catch up on lost sleep whenever weekday schedules are particularly hectic. It is known that women outperform men in academics, and they may enjoy a greater cognitive buffer against sleep loss.^27^ Tsai and Li conducted a sleep study on Taiwanese students and stratified some of the data by weekend versus weekday sleep.^28^ They reported a stronger correlation between sleep quality, rise time, time in bed, and sleep efficiency in men than women. Although they did not perform correlation analyses between sleep and GPA, they did report that women complained of more sleep difficulties on weekends, which may suggest that women cannot reap the same benefits as men from recovery sleep on the weekend. We, on the other hand, did not observe significant gender differences in sleep measures, such as time to fall asleep, during either the week or weekend.

We did not expect to find that weekday sleep would not be correlated with academic performance, because Okano and colleagues showed that sleep measured for the weeks before an exam (*i.e.*, during the learning phase) was indeed correlated with grades (the authors did not distinguish weekday from weekend sleep). The correlation Okano *et al.* observed between sleep duration and sleep quality was higher in men than women, and the authors wrote, “it may be more important for males to get a long-duration sleep in order to get good quality sleep.” Notably, Okano reported that sleep inconsistency was inversely correlated with grades for men but not women, and concluded, “it is important for males to stick to a regular sleep schedule in order to perform well in academic performance but less so for females.”

One explanation for the discrepancies between our work and that of others is, of course, differences in methodology: 1) We did not rely on self-reported GPAs, as in the work of Zeek *et al.*^22^ and Cates *et al.*^23^ Rather, we collected permission to acquire verifiable GPAs from the Dean’s office, which were sent there directly from the Duquesne University registrar. 2) We stratified sleep data into weekday versus weekend, unlike the sleep studies reported above, with the exception of the work of Tsai and Li.^28^ 3) The most obvious and important limitation of the current study was the reliance on self-reported sleep duration, including time to fall asleep (due to lack of financial resources), rather than wrist sleep monitors or electroencephalograms/polysomnography. Our sleep data were collected through a mandatory homework assignment on a daily basis for three weeks, and are therefore independent of lapses in long-term memory recall, which can compromise survey data integrity. Surveys are, however, vulnerable to any potential impact of gender on self-reporting. Future work on our student body with the Pittsburgh Sleep Quality Index, as in Cates *et al*,^23^ would be of additional value.

In the School of Pharmacy program at Duquesne University, exams in the first year of the professional phase are typically held outside of class time—early in the morning—before the large lecture halls are used for other courses. Two large lecture halls are required for each exam, as students are spaced far apart to lower the risk of cheating. Changing exam times to later in the day, after lectures of other classes have been completed and the large classrooms are free again, has met with resistance among our student body. Using a 5-point Likert scale, we asked the present cohort whether they wished to keep exams early in the day. In response, 53% of students stated that they preferred (*i.e.*, chose 4 or 5 on the Likert scale of agreement) to keep exams “early in the day, including at 07:30 AM”. These observations support anecdotal comments that the majority of students prefer to keep the evenings free for social activities, job-related work, and/or prefer not to be distracted by the looming presence of an upcoming exam for the earlier part of the day.

Nightly sleep durations considerably shorter or longer than 7 hours have been correlated with morbidity and the acceleration of death in a U-shaped pattern.^29^ This observation and a large body of work on the preponderance of U-shaped dose-response curves in biology^30^ might be construed to assume that long sleep durations *cause* poor health. However, those who are sick also tend to sleep more, and regularly forcing the infirm awake in an attempt to shorten their sleep durations to seven hours/night might worsen their health status and hasten an early death, as suggested by studies of intensive care unit delirium, the risk of which is reduced by the simple measure of using earplugs.^31^

In conclusion, we speculate that setting an early alarm on weekends in an effort to forcibly maintain the same sleep schedule as during the week may be counterproductive in students sitting for frequent early examinations or classes. To maintain a longer sleep duration, eye masks to block light transmission and earplugs might be helpful for students living in our college dormitories. Under ideal circumstances, however, students would go to bed earlier and maintain consistent sleep habits that keep their circadian timing systems properly aligned with the rest/activity rhythm.

## Acknowledgements

RKL conceived the study, wrote the paper, interpreted and analyzed data, and constructed the final figures. SLW entered and analyzed all the data, constructed figures, and contributed to experimental design, interpretation, and manuscript editing. MNC contributed to experimental design, collected survey data, and edited the manuscript. DCR contributed to experimental design and interpretation and edited the manuscript. We are indebted to the School of Pharmacy for their generous support of Dr. Leak’s lectures on the epidemiology and biological impact of sleep. We are also grateful to the Duquesne pharmacy students, for their kind participation. The authors have no conflicts to declare.

