## Appendix 1 for "Long weekend sleep is linked to stronger academic performance in male but not female pharmacy students"

### Survey One

(via Survey Monkey)

Question 1: What time did you go to bed?

Question 2: What time did you fall asleep?

Question 3: How many times did you wake up in the middle of the night? Why did you wake up?

Question 4: What time did you wake up in the morning?

Question 5: Did you feel refreshed within 30 minutes of waking up?

### Survey 2

ANONYMOUS QUESTIONNAIRE

*Please do not add personal identifiers; this evaluation is entirely anonymous.*

Please circle one of the following choices

1. I often study late at night even though I am tired and it is hard to concentrate

   Strongly disagree ------------------------------------------------------ Strongly agree

   1 2 3 4 5

2. I can most readily understand difficult material at these times of day:

   Mornings (8 AM to noon)

   Afternoons (noon to 4 PM)

   Evenings (4 PM to 8 PM)

   Early Nighttime (8 PM to midnight)

   Late Nighttime (midnight to 4 AM)

3. During the week, I typically sleep the following hours if there is an exam the next day:

   4-5 5-6 6-7 7-8 8-9 9-10 10-11 11+

4. During the week, I typically sleep the following hours if I ***don't*** have any exam the next day:

   4-5 5-6 6-7 7-8 8-9 9-10 10-11 11+

5. During the weekend, if I don’t have to get up early, I typically sleep the following hours:

   4-5 5-6 6-7 7-8 8-9 9-10 10-11 11+

6. Please add a comment about whether your sleep feels refreshing or you feel it is not sufficient:

7. My sleep is often interrupted by roommates or other sources of noise

Strongly disagree ---------------------------------------------------------- Strongly agree

1 2 3 4 5

8. My current GPA is

2.5 or less 2.51 – 3.0 3.01 – 3.5 3.51 – 3.70 3.71 – 4.0

9. My grade in Dr. Meng's exam in Human Physiology and Pathology was the following:

0-60% 61-70% 71-80% 81-90% 91-100%

10. My grade in Dr. Leak's exam in Human Physiology and Pathology was the following:

0-60% 61-70% 71-80% 81-90% 91-100%

11. My age is

21 or below 22 23 24 25 or older

12. My gender is __________________

13. I commute to school: Yes No (If the answer is "no," skip to question 15)

14. My commute time is, on average, about __________________ minutes

15. I prefer exams early, such as at 7:30 AM, rather than later in the day after other classes

Strongly disagree ---------------------------------------------------------- Strongly agree

1 2 3 4 5

This survey was stapled to a long consent form (available upon request) and then detached and stored separately to deidentify all the data

### Survey 3

1. Are you a transfer student? Yes No

2. Are you a commuter? Yes No

If yes, do you live: Alone With parents/family With friends With significant other

3. The area I live in is very noisy at night

Strongly disagree -----------------------------------------------------Strongly agree

1 2 3 4 5

4. Do you have roommates? Yes No

5. How many hours a day do you spend studying/doing homework when you don't have an exam?

_________________ hours

6. How many hours a day do you spend studying/doing homework when you do have an exam?

__________________hours

7. How many days a week do you take naps? ___________days

8. When you take a nap, how long do you typically nap? _________minutes

9. I usually feel refreshed after my nap

Strongly disagree -----------------------------------------------------Strongly agree

1 2 3 4 5

10. How many hours a week do you work? __________

11. How many credits are taking this semester? _________________

12. Are you on an athletic team? ____________________

11. Do you agree to allow Dr. Leak access to your GPA and grades in other classes?

Yes No

Signature: __________________________________________
